# Interactions between the 2C protein of FMDV and components of the viral replication machinery are mediated by ER-derived membranes

**DOI:** 10.1101/2025.09.17.676763

**Authors:** Connor Hayward, Eero V. Hietanen, Samuel J. Dobson, Morgan R. Herod, Juan Fontana, David J. Rowlands, Nicola J. Stonehouse

## Abstract

Foot-and-mouth disease virus (FMDV) remains an ever-present threat to the economic stability of the livestock industry and global trade. Despite this, questions remain regarding the fundamental biology underpinning the replication of this virus. Here, we examine components of the FMDV replication machinery (focussing on the viral 2C protein) and investigate the conditions under which they interact. Using a novel 2C antibody in co-immunoprecipitation experiments under different conditions followed by mass spectrometry, we identify membrane-associated proteins (such as the viral proteins 2B and precursors of 3A, which are poorly-characterised proteins involved in viral replication) along with ER-associated host proteins. In addition, our analysis shows that a number of nuclear factors interact with 2C in a membrane-independent manner, potentially being co-opted to support RNA replication of the virus. Furthermore, we demonstrate that the interaction of 2C with several viral proteins (including key members of the replication machinery and viral RNA) is maintained following ultracentrifugation, suggesting that these co-sediment as part of a complex. Our data suggest that the replication complex is ER-derived and highlight several new avenues of investigation for the disruption of the FMDV lifecycle.

## Introduction

Foot-and-mouth disease (FMD) is an animal disease of global concern affecting food security and commerce. Foot-and-mouth disease virus (FMDV), the causative agent of FMD, remains endemic across many parts of the globe, posing a constant threat to FMD-free nations. FMD affects cloven-hoofed animals, including major livestock species, and losses associated with the disease (and subsequent control measures) can be economically damaging (Fenner, 2017). For example, the 2001 United Kingdom (UK) outbreak cost an estimated >£13 billion and resulted in the slaughter of >4 million animals (National audit office, 2002). FMD also impacts those reliant on subsistence farming in endemic regions and has been linked to malnutrition in children by lost access to milk produced by livestock (Bayissa *et al*., 2011; Barasa *et al*., 2008; Knight-Jones, McLaws and Rushton, 2017). Recent outbreaks have occurred in Germany, Hungary and Slovakia in 2025, highlighting the ongoing concerns posed by FMDV.

The organisation of the FMDV RNA genome is similar to that of other picornaviruses, with untranslated regions flanking the single open reading frame (ORF). The ORF is initially processed into P1/2A (the structural proteins, VP1-4), P2 and P3 (non-structural proteins), through the action of viral proteases and ribosomal skipping (Grubman and Baxt, 2004; Ryan, Belsham and King, 1989; Grubman *et al*., 1984). Further processing of these polyproteins by the virally encoded protease 3C (and its precursor 3CD) results in production of 14 individual proteins, including the RNA-dependent RNA polymerase (3D), the RNA replication priming peptides (3B_1-3_) and the less well-characterised 2C and 3A proteins. Polyprotein processing also produces a cascade of precursors, some of which have independent roles in replication e.g. 3CD (Herod *et al*., 2017; Pierce *et al*., 2023).

FMDV genome replication occurs in cytoplasmic membranous complexes (Monaghan *et al*., 2004), but less is known about FMDV replication complexes than those of the related picornavirus, poliovirus (PV). However, significant differences have been reported. Firstly, FMDV replication is insensitive to brefeldin A, an inhibitor of endoplasmic reticulum to Golgi trafficking using COP-I, in stark contrast to PV (Gazina *et al*., 2002; Martín-Acebes *et al*., 2008; Midgley *et al*., 2013). Secondly, FMDV replication is not dependent on the activity of phosphatidylinositol 4-kinase (PI4K) and no co-localisation has been observed between PI4P lipids (which are produced as a result of this activity) and non-structural viral proteins. Additionally, the level of PI4P does not increase during infection and knockdown of PI4K has no effect on viral replication (Berryman *et al*., 2012; Loundras *et al*., 2016; Berryman *et al*., 2016). Thirdly, the role of PV 3A in disrupting ER-Golgi trafficking appears to be substituted in FMDV by 2BC, since expression of FMDV 3A alone does not impact this pathway, in contrast to expression of PV 3A (Moffat *et al*., 2005; Moffat *et al*., 2007). While generalities of FMDV replication can be inferred from studies of other picornaviruses, these three examples illustrate that not all features are shared across picornaviruses.

In order to understand the biogenesis and morphology of FMDV replication complexes multiple studies have used immunofluorescence to probe for co-localisation of FMDV proteins with host markers of cellular organelles. However, such associations are elusive (Knox *et al*., 2005; Berryman *et al*., 2012; Berryman *et al*., 2016) and the origin of FMDV replication complex membranes remains unclear. Fluorescence microscopy studies have demonstrated co-localisation of FMDV 3A with pre-Golgi membranes in a peri-nuclear manner, as has been observed in early PV infection, where ER membranes are implicated (Midgley *et al*., 2013; Melia *et al*., 2019). However, electron microscopy studies suggest that the ultrastructure of FMDV replication complexes differs from those of PV (Monaghan *et al*., 2004). FMDV induces numerous small single membrane vesicles that accumulate in a perinuclear distribution during replication. This is in contrast to PV, which despite sharing a similar sub-cellular distribution, forms rosette-like membrane clusters when visualized using similar resin embedding and sectioning techniques (Monaghan *et al*., 2004; Bienz *et al*., 1980). More recent cryo-EM imaging of PV replication complexes has suggested that these rosette arrangements may be more complex tubular networks, something which has yet to be interrogated for FMDV (Dahmane *et al*., 2022). Taken together, these observations suggest that FMDV replication complexes may differ structurally from those of the better characterised replication complexes of other picornaviruses and may involve different factors/cellular components.

While picornaviral 2C has been extensively characterised biochemically, the exact function and role performed by this protein during replication remains unclear (Sweeney *et al*., 2010; Zhang *et al*., 2022). Guanidine hydrochloride (GnHCl) is a potent inhibitor of the replication of many positive sense RNA viruses, including FMDV, and resistance to the compound is associated with the acquisition of mutations in the 2C protein, underscoring its importance to replication (Pincus and Wimmer, 1986; Sweeney *et al*., 2010; Saunders *et al*., 1985). Picornavirus 2C proteins have ATPase activity and bind ssRNA non-specifically. There is growing evidence for the formation of multimeric structures by 2C-like proteins (Zhang *et al*., 2022; Haas *et al*., 2025). However, despite their inclusion in the helicase super-family, helicase activity has been rarely reported (Sweeney *et al*., 2010; Xia *et al*., 2015). Recent structural evidence from cryo-EM tomography, as well as previous genetic evidence, has suggested that 2C of PV may tether viral capsids to the surface of replication complexes during virus assembly (Dahmane *et al*., 2022; Liu *et al*., 2010). Whether this is a function shared by 2C proteins of other picornaviruses remains to be demonstrated, but this does suggest that 2C is a key protein in replication. FMDV 2C has been shown to co-localise with other viral non-structural proteins, such as 3A and 3D, at putative sites of replication in the perinuclear space (Knox *et al*., 2005). However co-localisation of FMDV 2C with host-cell organelle marker proteins has yet to be demonstrated, (Knox *et al*., 2005; Moffat *et al*., 2005; Moffat *et al*., 2007; Berryman *et al*., 2012; Midgley *et al*., 2013).

Questions therefore remain over the utilisation of host membranes for the formation of FMDV replication complexes and their composition. Here, we have elucidated the relationship between 2C and other viral proteins with host membranes. Specifically, we present evidence of ER membrane-mediated interactions between 2C and several other viral proteins, including 3A and 3D and their precursors. Additionally, we demonstrate that 2C, 3A, 3D and other proteins present in these interactions co-purify with viral RNA and provide evidence for the involvement of the ER in localisation of FMDV replication machinery and complex formation.

## Methods

### Cell lines

BHK-21 cells obtained from ATCC (LGC Standard) were maintained in Dulbecco’s modified Eagle’s medium (DMEM) with glutamine (Sigma-Aldrich) supplemented with 10% FCS, 50 U/mL penicillin, and 50 μg/mL streptomycin.

### *In vitro* transcription

The construction, linearisation and purification of plasmids containing wild-type FMDV replicons pRep-ptGFP and pRep-ptGFP-GNN were described previously (Tulloch *et al*., 2014; Herod *et al*., 2016). Linear DNA was transcribed *in vitro* using T7 RiboMAX Express Large Scale RNA Production System (Promega) in half-sized reactions with 250 ng input DNA. Reactions were incubated for 1.5 hours at 37°C before addition of 1 μL DNase 1 followed by a further 20 min incubation at the same temperature. The RNA produced was column purified using RNA Clean and Concentrator (Zymo Research), quantified by NanoDrop (Thermo Fisher) and integrity confirmed by denaturing MOPS-formaldehyde gel electrophoresis.

### Immunofluorescence

Transfection of FMDV replicons into BHK-21 cells has been described previously (Tulloch *et al*., 2014; Herod *et al*., 2016). Briefly, BHK-21 cells were seeded onto coverslips before transfection with replicon RNA using Lipofectamine 2000 (Thermo Fisher) at a ratio of 3 μL:1μg (reagent:RNA). Samples were fixed using 4% paraformaldehyde at desired time-points post-transfection and subsequently washed with PBS. Primary antibodies used were rabbit anti-2C 3-2 (raised against 2C peptide; VEMKRMQQDMFKPQP, by Custom Polyclonal Antibody Services) and mouse anti-3A 2C2 (a kind gift from Francisco Sobrino). Secondary antibodies used for detection were anti-rabbit Alexa fluor 555 and anti-mouse Alexa fluor 647 (Life Technologies). Cells were permeabilised with 0.1% Triton X-100, washed in PBS three times and blocked in PBS containing 1% BSA for 1 hour. Antibodies were diluted in blocking buffer with primary incubation at 37°C for 1 hour and secondary at room temperature for 30 min. Coverslips were mounted using ProLong Glass containing NucBlue (Thermo Fisher). Samples were visualised using a Zeiss LSM880 confocal microscope and processed using Zen Blue Software (Zeiss).

### Harvest and homogenisation of cells

Cell homogenates for immunoprecipitation and gradient purification were prepared in the same manner and scaled for material requirements. Cells were seeded onto dishes of 100 mm in diameter and transfected with replicon RNA as described previously, scaled as appropriate (Tulloch *et al*., 2014; Herod *et al*., 2016). Cells were harvested at indicated time-points post-transfection by scrapping and subsequently pelleted by centrifugation at 500 x *g* for 5 min. Pellets were washed twice in PBS, before being resuspended in 1 mL of homogenisation buffer (HB; 250 mM sucrose, 3 mM imidazole [pH 7.4], 2 µg/mL actinomycin D (Sigma), Pierce EDTA-free protease inhibitor (1 tablet per mL; Roche), 1 µL/mL RNaseOUT™ (Thermo Fisher), pelleted and resuspended again in 500 µL of fresh HB. Homogenisation was performed using a 22-gauge needle with 24 strokes on ice. Samples were centrifuged at 500 x *g* for 10 min at 4°C to pellet nuclei and non-lysed cells. Supernatant (post-nuclear supernatant (PNS)) was removed and transferred to fresh tubes.

### Immunoprecipitation

Non-specific binding proteins were removed from PNS (200 µL) by incubating with 20 µL of non-conjugated Protein A DynaBeads (Thermo Fisher) for 15 min at room temperature on a rotating mixer, following which the beads were removed by magnetic separation. Protein A DynaBeads (Thermo Fisher) prewashed in IP wash buffer (10 mM Tris pH 7.4, 1 mM EDTA, 150 mM NaCl) were mixed with anti-2C 3-2 rabbit antibody at a ratio of 2:1 and incubated for 1 hour at room temperature on a rotating mixer. Remaining excess non-conjugated antibody was removed by three 15 min washes with IP wash buffer. Conjugated beads were then mixed with pre-adsorbed PNS at a ratio of 1:1 volume to volume (in IP wash buffer) and rotated at room temperature for 1 hour. Antibody-bound complexes were purified by magnetic separation and the beads were washed three times for 15 min using IP wash buffer. Proteins were eluted from the beads using 150 µL of 0.2 M glycine pH 2 in IP wash buffer. Eluate was neutralised by addition of 150 µL pH 8.5 Tris-base in IP wash buffer and stored at -20 °C.

To examine the susceptibility of protein complexes to detergent, different concentrations of Triton X-100 were added to pre-adsorbed PNS prior to mixing with anti-2C conjugated beads and rotating at room temperature for 30 min. To examine the effects of salt on binding, NaCl in HB at twice the final indicated concentration was mixed 1:1 with PNS for 30 min with rotating at room temperature. Treatment of PNS with RNase A was performed by the addition of 1 unit/µL of PNS and incubating at room temperature for 1 hour while rotating. All pre-treatment experiments then followed the methods described above.

### Mass spectrometry

Samples from immunoprecipitation were analysed on bead by tandem mass tag mass spectrometry at the University of Bristol proteomics facility. Raw data was extracted using Protome Discoverer (Thermo Fisher Scientific). The vendor .raw format files were subsequently converted to .mzML files using MSConvert v3.0 (Chambers *et al*., 2012) to permit downstream analysis using FragPipe v22.0 (Kong *et al*., 2017; Yang *et al*., 2023; Teo *et al*., 2021; da Veiga Leprevost *et al*., 2020; Yu, Haynes and Nesvizhskii, 2021). Database search and quantification were performed against the UniProt *Cricetulus griseus* (Chinese hamster, taxonomic ID 10029) reference proteome (proteome ID UP000001075) combined with the full FMDV reference genome sequence including the mature and precursor protein sequences of FMDV P2 and P3 regions. Full search database consisted of 48,050 sequences including decoys. The standard TMT16-MS3 workflow available in FragPipe was used for the analysis, with the added modification of enabling MSBooster to improve peptide-to-spectrum scoring.

Background binding was accounted for through the filtering out of proteins identified as having log_2_ fold change between -0.5 and 0.5 in pairwise comparisons of wild-type versus mock for both non-treated and detergent treated conditions. Remaining proteins were analysed pairwise between wild-type samples non-treated versus detergent treated. A custom R script (R v4.4.1) was developed to perform downstream analysis after the database search. The R package MSstatsTMT v2.12.1 (Huang *et al*., 2020) was used to detect differences in protein abundances. Further enrichment analysis was performed using clusterProfiler v4.12.6 (Xu *et al*., 2024). Gene Set Enrichment Analysis (GSEA) using clusterProfiler was done by ranking the data based on the log_2_ fold-change and building GO term and protein ID association tables. Additional GO annotation sets for *Cricetulus griseus* were also fetched using QuickGO (Binns *et al*., 2009) from the UniProt annotation entries to explore the enrichment associated with ER, endoplasmic membrane, and endoplasmic lumen GO terms. The GSEA analysis results were then visualised as ridgeline plots for the experimental condition. Visualisation of the results and plotting was performed using the EnhancedVolcano (Blighe K, 2025), ggplot2 (H, 2016) and ggridges packages, while general data handling utilised the tidyverse package (Wickham H, 2019).

### Western blotting

Immunoblotting was performed as previously described (Forrest *et al*., 2014). Primary antibodies used were rabbit anti-2C 3-2 (Custom polyclonal antibody services), mouse anti-3A 2C2 and rabbit anti-3D 397 (both kind gifts from Franciso Sobrino). A mouse anti-GAPDH (Invitrogen; GA1R) primary antibody was used as a loading control. Appropriate species secondary antibodies, anti-rabbit DyLight™ 800 and anti-mouse DyLight™ 680 (Thermo Scientific) were used for detection and blots visualised using a LiCor Odyssey SA imager (LiCor). Densitometry was performed using ImageStudio software (LiCor).

### Primary gradient purification

PNS was further processed by isopycnic purification using discontinuous iodixanol gradients. Primary gradients (17 mL) were made by the underlaying of increasing percentage of iodixanol (10 to 30%) (Thermo Fisher) in dilution buffer (DB; 250 mM sucrose, 35 mM HEPES [pH 7.4], 7 mM KCl, 2.5 mM DTT, Pierce EDTA-free protease inhibitor (1 tablet per 30 mL; Roche) 1 µL/mL RNase Out (Thermo Fisher).

PNS was diluted 1:1 with HB to give a sample volume of 1 mL which was overlaid on gradients. Primary gradients were centrifuged at 100,000 x *g* for 3 hours at 4°C using a SW-32 Ti rotor in an Optima XPN-80 ultracentrifuge (Beckman Coulter). Following centrifugation, 1 mL fractions were taken, and samples analysed by SDS-PAGE or scintillation counting. Where fractions were taken forward for secondary gradient, 500 µL of each was pooled to form a 1.5 mL sample.

### Secondary gradient purification

Secondary gradient purification was performed on 1.5 mL of pooled primary gradient fraction sample. The sample was mixed with iodixanol to give a final percentage of 40% iodixanol. This was then underlaid below the secondary iodixanol gradient (17 mL, 10 to 30%). Secondary gradients were centrifuged at 90,000 x *g* for 18 hours at 4°C using a SW-32 Ti rotor in an Optima XPN-80 ultracentrifuge (Beckman Coulter). Following centrifugation, gradients were fractionated in 1 mL fractions, which were analysed by SDS-PAGE or scintillation.

### Tritiated uridine incorporation and scintillation

Cells were seeded on 100 mm diameter dishes as previously described and pre-treated for 1 hour with 10 μg/mL actinomycin D diluted in cell culture medium. Following pre-treatment, media was replaced with fresh media and transfection performed as described above. Radiolabelling of nascent RNA was performed by the addition of 500 μCi [^3^H] uridine (Hartmann) to each dish. At 4 hours post transfection cells were harvested, homogenized and gradient-purified as described above. Scintillation was performed by mixing 500 μl of each gradient fraction with 3 mL of scintillation fluid (Perkin Elmer). Radioactive decay from samples was counted for one minute using a TriCarb scintillation counter (Perkin Elmer) and background subtraction was performed using scintillation fluid mixed with non-radiolabelled fraction sample.

## Results

### Characterisation of 2C expression and localisation using a novel antibody

Previous studies have shown both 2C and 3A proteins to be essential for FMDV replication (Sweeney *et al*., 2010; Gao, Sun and Guo, 2016) and several have also shown a close association between 3A and putative replication complexes (Moffat *et al*., 2005; Moffat *et al*., 2007). Due to a lack of available reagents to study FMDV 2C, we generated a novel antibody against a peptide of FMDV 2C (residues highlighted on the previously published crystal structure of FMDV 2C, Figure S1.A). In order to characterise this novel antibody, we performed an immunofluorescence and western blot time-course in comparison with an anti-3A antibody (Seago *et al*., 2012). Experiments were performed using a FMDV replicon model in which the viral structural proteins have been replaced with a fluorescent marker protein, ptGFP, thus enabling replication to occur in the absence of virion production (Figure S1.B) (Tulloch *et al*., 2014; Herod *et al*., 2015; Loundras *et al*., 2016).

Cell lysates were harvested at indicated times post-transfection with the replicon and subjected to western blot analysis. FMDV proteins exist as both precursor forms and as fully cleaved mature proteins. Antibodies capable of binding the mature forms of proteins will likely also bind their precursors. Precursors containing 2C and/or 3A can be separated by mass and charge during SDS-PAGE and can be identified by immunostaining with anti-2C and anti-3A antibodies. GAPDH was probed across all time-points as a loading control. As expected, densitometry showed decreases in large precursors over time (e.g. 3AB_1-3_CD) with concomitant increases in mature 2C and 3A, which were maximal at 4-to 5-hours post transfection (hpt) (Figure 1.A, Figure S2). A number of 3A precursors linked to 3B peptides were also detectable from 2-hpt, some appearing as doublets, possibly due to post-translation modifications. The detection of 3A precursors has been reported previously using the anti-3A (2C2) antibody (Seago *et al*., 2012). For consistency, densitometry was performed using the lower band of any doublet related to the 3A precursors.

**Figure 1.**
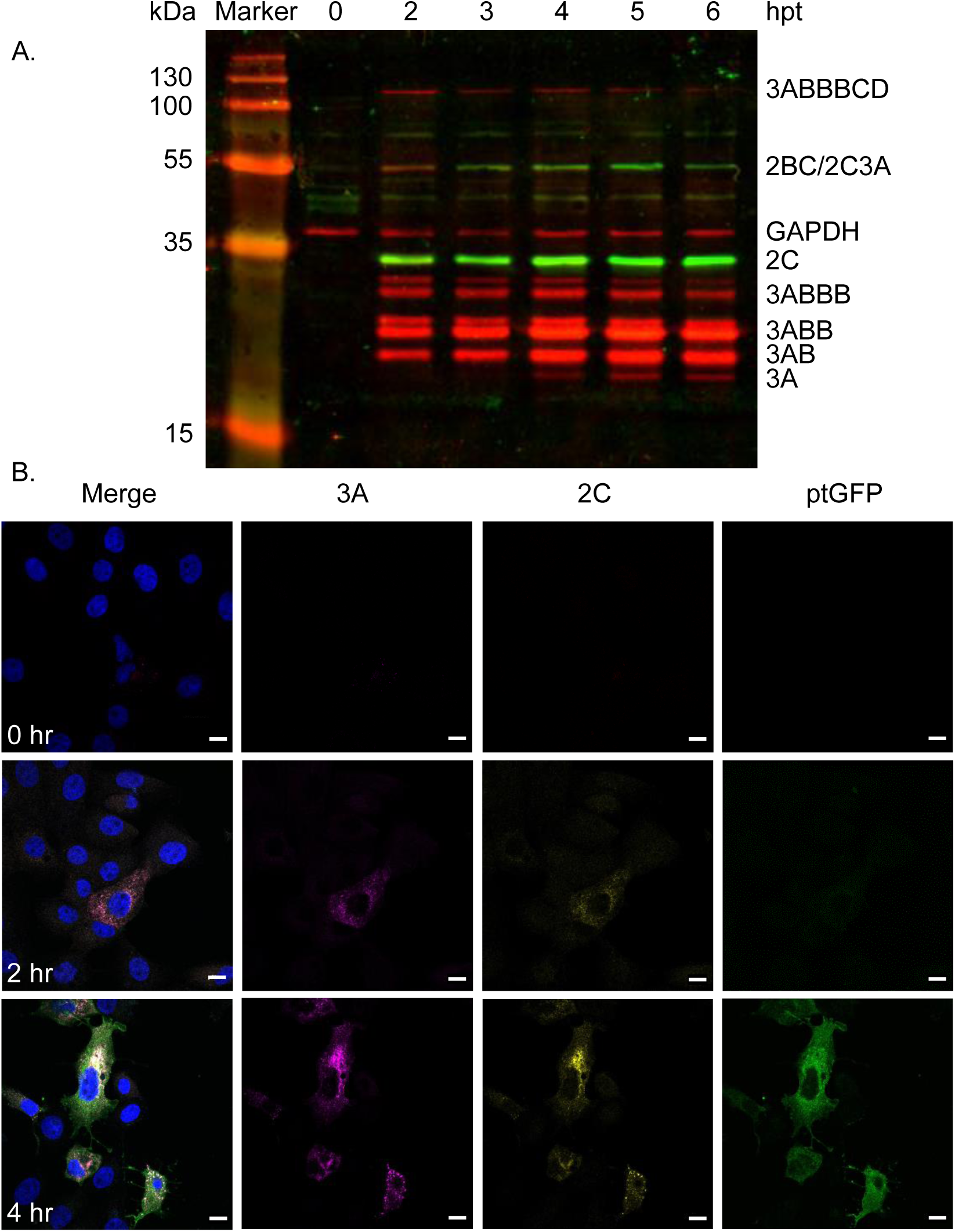
Western blot and immunofluorescent analyses of a FMDV replicon using a novel 2C antibody. (A) BHK-21 cells were transfected with FMDV replicon and harvested, before protein separation by SDS-PAGE and subsequent western blotting with anti-2C (green), anti-3A (red) and GAPDH (red) followed by DyLight™ 680 and DyLight™ 800 secondary antibodies. Mature and precursor proteins are labelled according to expected molecular weights. (B) Confocal microscopy of anti-2C, anti-3A and ptGFP in BHK-21 cells fixed at indicated times post-transfection. Cells were immunostained with anti-2C followed by anti-rabbit Alexafluor-647; and anti-3A followed by anti-mouse Alexafluor-555. Samples were stained with DAPI and visualised using a Zeiss LSM880 confocal microscope. Scale bars = 10 µm.

Co-localisation of 2C and 3A proteins (and potentially also their precursor forms) was expected to occur during replication and was subsequently demonstrated by indirect immunofluorescence from 2-hpt (Figure 1.B). The signals for both antibodies were cytoplasmic, predominantly around the perinuclear region with puncta visible in some cells (Figure 1.B). ptGFP fluorescence was detected later, possibly due to its diffuse distribution throughout the cytoplasm (Figure 1.B). Having demonstrated that our anti-2C peptide antibody appeared to have low non-specific activity and to co-localise with an established and previously published anti-3A antibody, we used the 2C reagent to further investigate the role of the 2C protein in FMDV replication using the replicon model.

### Complexes of 2C with 3A and 3D precursor proteins are detergent sensitive

Immunoprecipitation of transfected cell lysates was used to assess potential interactions between 2C and other viral proteins. Performing this with anti-2C on lysates from cells 4-hpt with wild-type replicon yielded a robust signal for 2C, as expected (Figure 2). Probing the anti-2C bound material with anti-3A and anti-3D revealed strong signals for precursors at the expected sizes of 3AB and 3ABB (and possibly 3ABBB) as well as 3CD. Incubation of lysates in a range of Triton X-100 concentrations prior to immunopreciptation showed that co-immunoprecipitation of 3AB, 3ABB and 3CD with 2C could be disrupted by 1% and 0.1% Triton X-100, but were unaffected by 0.01% detergent (Figure 2). Probing for mature 3D was not feasible following IP probably due to the cross-reactivtiy of secondary antibodies with the eluted heavy chains of the antibodies used for IP, thus masking the 55 kDa signal. However, overall these data suggested that interactions between these proteins and 2C are membrane-mediated.

**Figure 2.**
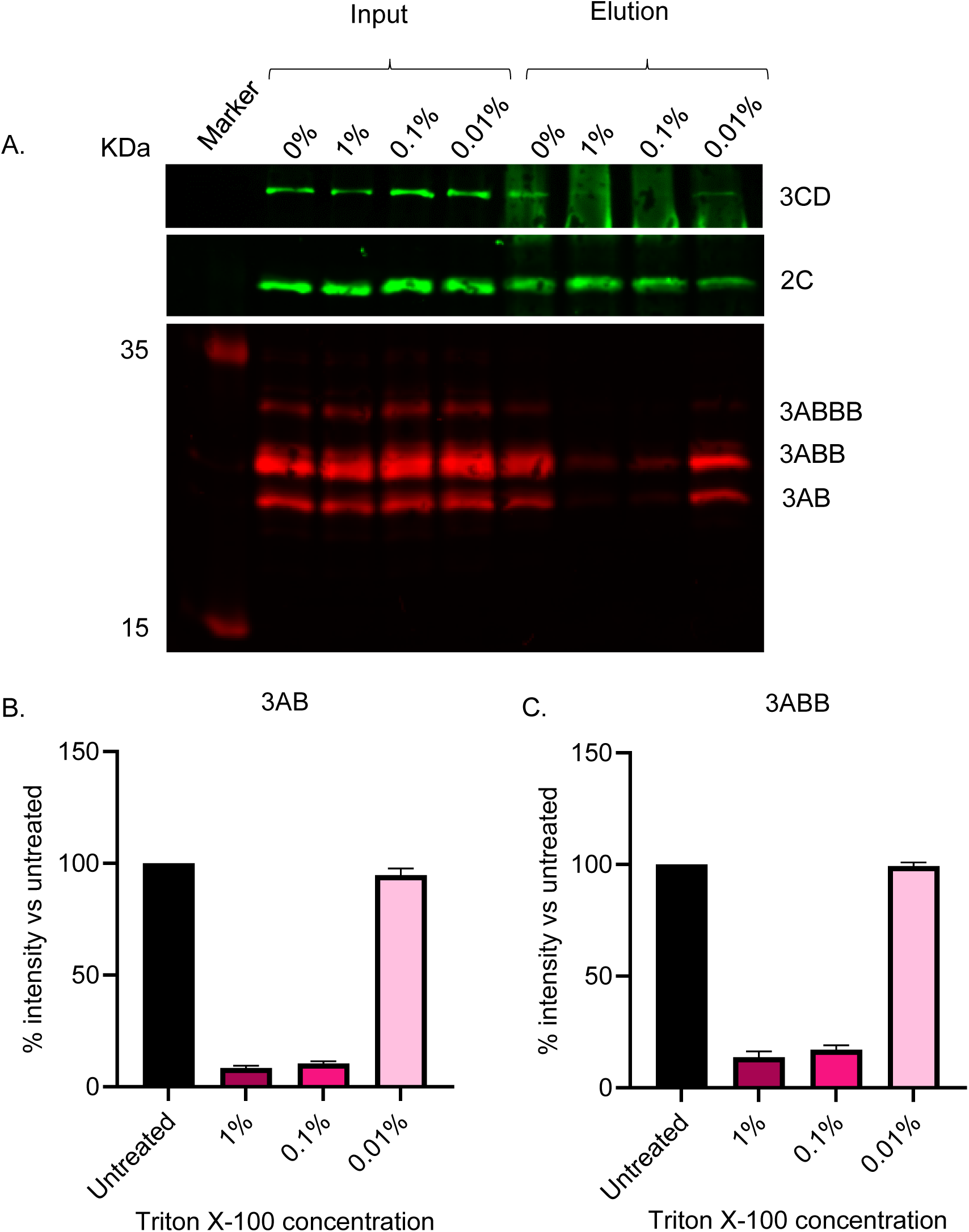
Western blot analysis following anti-2C immunoprecipitation from lysates of cells transfected with wild-type replicon and treated with Triton X-100. Cells were harvested 4-hpt with FMDV wild-type replicon. Lysates were subsequently treated with Triton X-100 (0%, 0.01%, 0.1% and 1%) for 30 minutes prior to anti-2C IP. (A) SDS-PAGE and western blot analyses was performed on input and eluate samples using primary antibodies against 3D (green), 2C (green) and 3A (red) followed by anti-mouse DyLight™ 680 and anti-rabbit DyLight™ 800 secondary antibodies. Densitometry was performed comparing signal intensity of 3AB (B) and 3ABB (C) with untreated samples. A is a representative result of 3 biological repeats. Error bars represent standard deviation (n = 3).

To further investigate the nature of the protein associations within the presumed complexes, lysates were treated with a range of NaCl concentrations prior to co-immunopreciptation with anti-2C. High salt concentrations can disrupt electrostatic interactions between proteins while leaving membrane mediated interactions unaffected. Again, 2C co-immunoprecipitated with 3A precusors and 3CD in control conditions and interactions with 3A precusors were maintained across salt concentrations (Figure 3). The 3CD signal decreased at 0.5 M and 1 M salt, although (as above) quantification was not reliable due to reactivity of the secondary antibody with the heavy chains of the antibodies used for IP (Figure 3). However, the insensitivity of 3A precusors co-immunoprecipiation with 2C to elevated salt concentrations further supports the hypothesis that these proteins interact in a membrane-mediated fashion.

**Figure 3.**
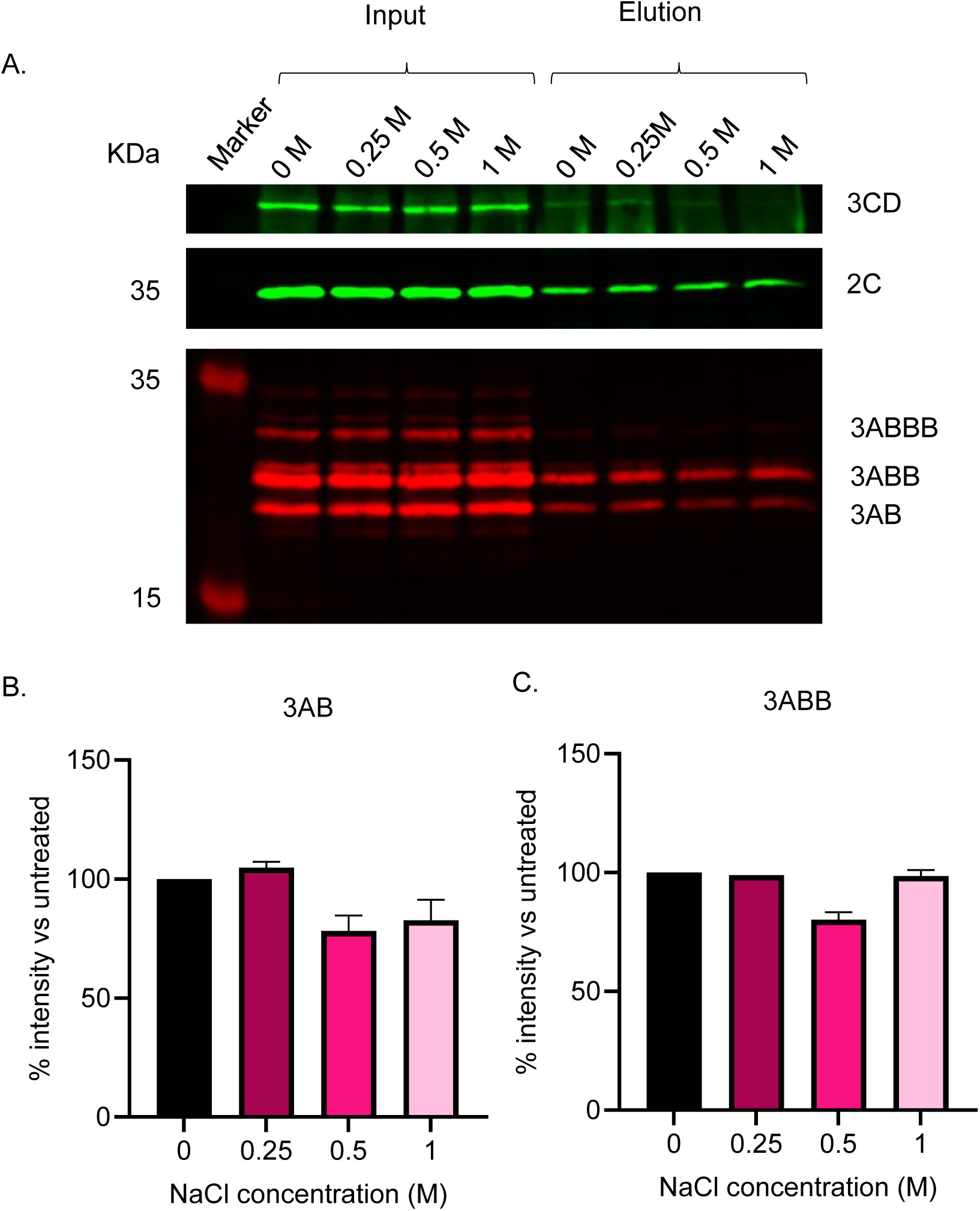
Western blot analysis following anti-2C precipitation of cell lysates transfected with wild-type replicon and treated with a range of NaCl concentrations. Cells were harvested 4-hpt with FMDV wild-type replicon. Lysates were subsequently treated with a range of NaCl (0 M, 0.25 M, 0.5 M and 1 M) for 30 minutes prior to anti-2C immunoprecipitation. (A) SDS-PAGE and western blot analysis was performed on input and eluate samples using primary antibodies against 3D, 2C and 3A with anti-mouse DyLight™ 680 and anti-rabbit DyLight™ 800 secondary antibodies. (B and C) Densitometry performed comparing signal intensity of 3AB (B) and 3ABB (C) to untreated samples. A is a representative result of 3 biological repeats. Error bars represent standard deviation (n = 3).

### Viral proteins and RNA co-sediment during ultracentrifugation

As 2C appears to interact with other viral proteins in a membrane-mediated fashion, attempts were made to purify 2C-containing membranous complexes. Cell lysates were separated on iodixanol density gradients followed by western blotting of fractions to identify viral proteins. To assist purification, a two-stage procedure was performed in which lysate was first overlayed on a primary discontinuous gradient of iodixanol, followed by the pooling of selected fractions which were then underlayed below a secondary discontinuous iodixanol gradient, as detailed below.

Western blotting with anti-2C, anti-3A and anti-3D across all gradient fractions was performed to detect mature proteins and precursors, as demonstrated previously. As expected, signals corresponding to 2C, multiple 3A precursors, 3D and 3CD were detected in the input sample. Within the primary gradient fractions, 2C and 3A precursors were present in a broad peak encompassing fractions 9-12, corresponding to ∼20% iodixanol (Figure 4.A). A similar distribution was observed for 3D and 3CD, although a larger proportion of signal remained at the top of the gradient in fractions 1 and 2 (Figure 4.A; Figure S4). Control gradients using material produced following transfection with GNN replicon yielded low level signals as would be expected from input translation (Figure S3).

**Figure 4.**
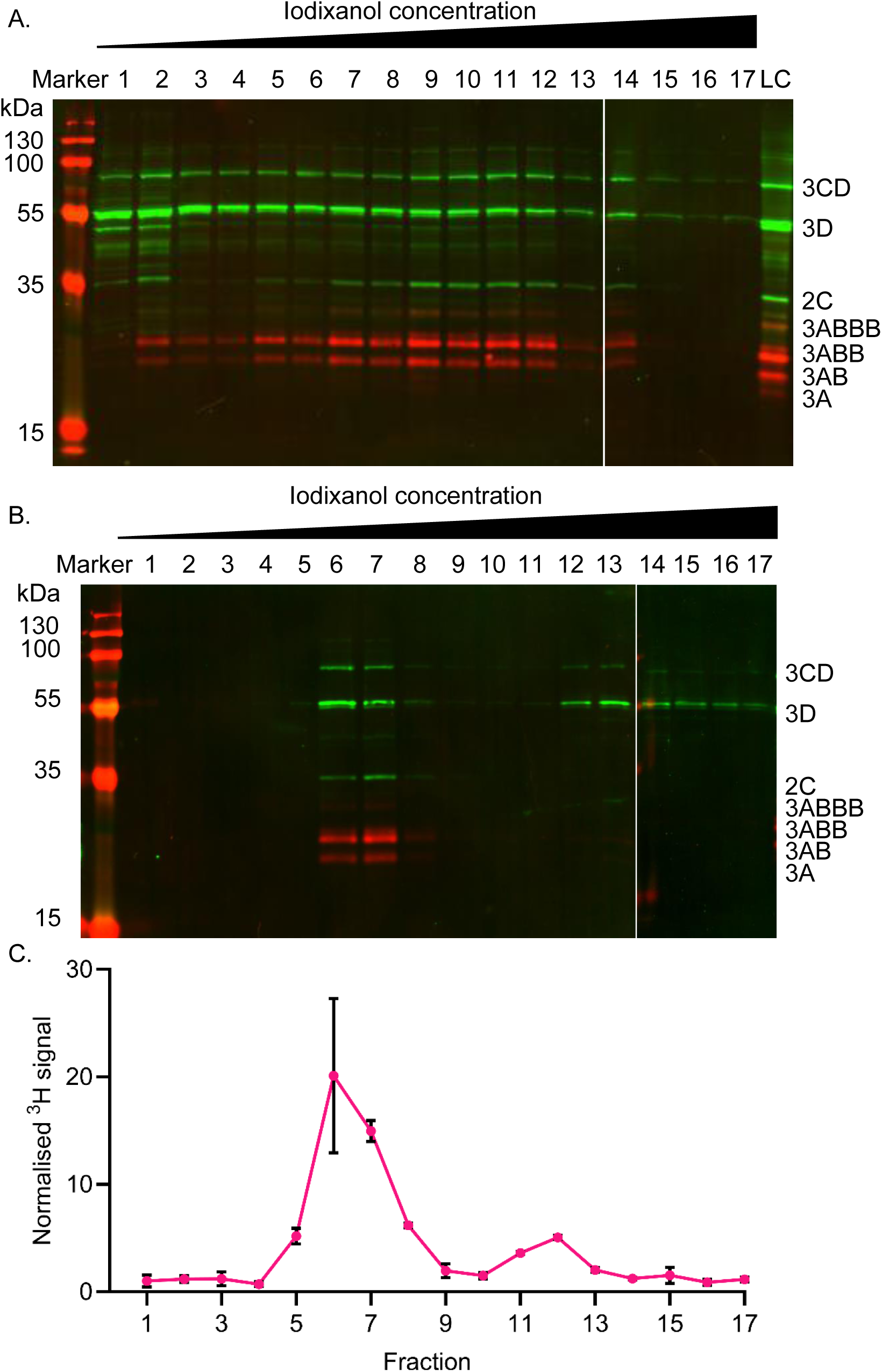
Co-sedimentation of FMDV non-structural proteins and RNA during density gradient ultracentrifugation. BHK-21 cells transfected with wild-type FMDV replicon were harvested 4-hpt and homogenised. (A) Lysate was overlaid on discontinuous iodixanol gradients (10%-30%) prior to ultracentrifugation. Proteins from each 1 mL fraction were separated by SDS-PAGE prior to western blotting with anti-2C, anti-3A and anti-3D antibodies. Anti-rabbit DyLight™ 800 (Green) and anti-mouse DyLight™ 680 (Red) secondary antibodies were used for detection of respective primary species. Precursor proteins are labelled based on expected molecular weights. A representative result of 3 biological repeats is shown. (B) Fractions 9-11 from each primary gradient were pooled and underlaid beneath a secondary discontinuous iodixanol gradient. Following ultracentrifugation, fractions were probed by western blot as above. A loading control (LC) of wild-type lysate was included for normalisation of densitometry across western blots. (C) Fractions 9-10 from tritiated primary gradients were pooled and subjected to the same secondary discontinuous iodixanol gradient (10-40%). Following ultracentrifugation, [^3^H] detection was undertaken by scintillation. [^3^H] signal was normalised relative to paired GNN control gradients. Error bars represent standard deviation (n = 3).

Having noted the co-sedimentation of 2C with multiple viral proteins, a potential indication of a replication complex, fractions corresponding to ∼20% iodixanol were pooled and subjected to further purification by underlaying below a secondary gradient. As expected, this additional ultracentrifugation step increased the proportions of 2C, 3AB and 3ABB signal in fractions 6 and 7, corresponding to ∼20% iodixanol, with little signal detected in other parts of the gradient. Similarly, accumulation of both 3D and 3CD was observed within fractions 6 and 7, although a proportion of each remained within the fractions corresponding to higher iodixanol concentrations suggesting a more transient or weaker association between these proteins and the remainder of the complex co-sedimenting 2C and 3A precursors (Figure 4.B; Figure S5).

Having established that the viral proteins previously shown to co-immunoprecipitate also co-sedimented, we investigated the presence of viral RNA as part of a replication machinery. In order to do this, cells were treated with actinomycin D to halt host RNA transcription and subsequently transfected with wild-type or GNN replicon RNA in the presence of [^3^H] uridine. The incorporation of [^3^H] uridine into the replicating viral genome is unaffected by actinomycin treatment (Black and Brown, 1969; Adeyemi *et al*., 2021). Lysates were then harvested and purified in the same manner as those for protein analysis and incorporation of [^3^H] uridine assessed by scintillation counting of the gradient fractions. To control for the migration of unincorporated [^3^H] uridine into the gradient, GNN replicon transfected lysates were purified in parallel and the signal used for normalisation of wild-type lysates. The profile of the normalised [^3^H] signal from wild-type transfected lysate was similar to that seen by western blotting. The peak [^3^H] signal was present within fractions 6-8 of secondary gradients corresponding to ∼20% iodixanol (Figure 4.C), similar to that of 2C and 3A previously observed (Figure 4.B) and suggesting the successful purification of FMDV replication machinery.

### Identification of anti-2C immunoprecipitated proteins by mass spectrometry following detergent treatment showed enrichment of ER proteins

Earlier studies have investigated the origin, biogenesis and composition of FMDV replication complexes using a variety of methodologies, yet the contents and origins of the membranes involved are still debated. Having demonstrated by co-immunoprecipitation the association of 2C with 3A and 3D, we probed for viral and host proteins using tandem mass tag (TMT) mass spectrometry on both non-treated and detergent-treated lysates, harvested from either mock or FMDV replicon transfected cells. In order to improve stringency and reliably when conducting differential enrichment analysis between non-treated and detergent-treated transfected lysates, peptide hits from mock treated samples which had log_2_ fold change (FC) between -0.5 to 0.5 were considered as non-specific antibody binding proteins and removed. Some potential 2C-interactors may therefore have been removed from the final analysis.

Following the removal of background from the datasets, differential enrichment analysis was performed between non-treated and detergent-treated transfected lysates. This was reflected in the output of the differential enrichment analysis, where peptides mapping to 3A and its detectable precursor 3AB_1_ displayed a relative log_2_ FC of 1.7 and 1.5, respectively (Figure 5.A). Using this approach, we were able to identify changes in another non-structural protein, 2B, which was previously not possible due to a lack of antibody reagents. The relative enrichment of 2.1 log_2_ FC of 2B following treatment with detergent, was even greater than that of 3A. Additionally, peptides mapping to 3D and 3CD were relatively unaffected by the presence of detergent with log_2_ FC of 0.2 and 0.3, respectively. As expected, 2C was unaltered as this was the direct target for immuno-precipitation.

**Figure 5.**
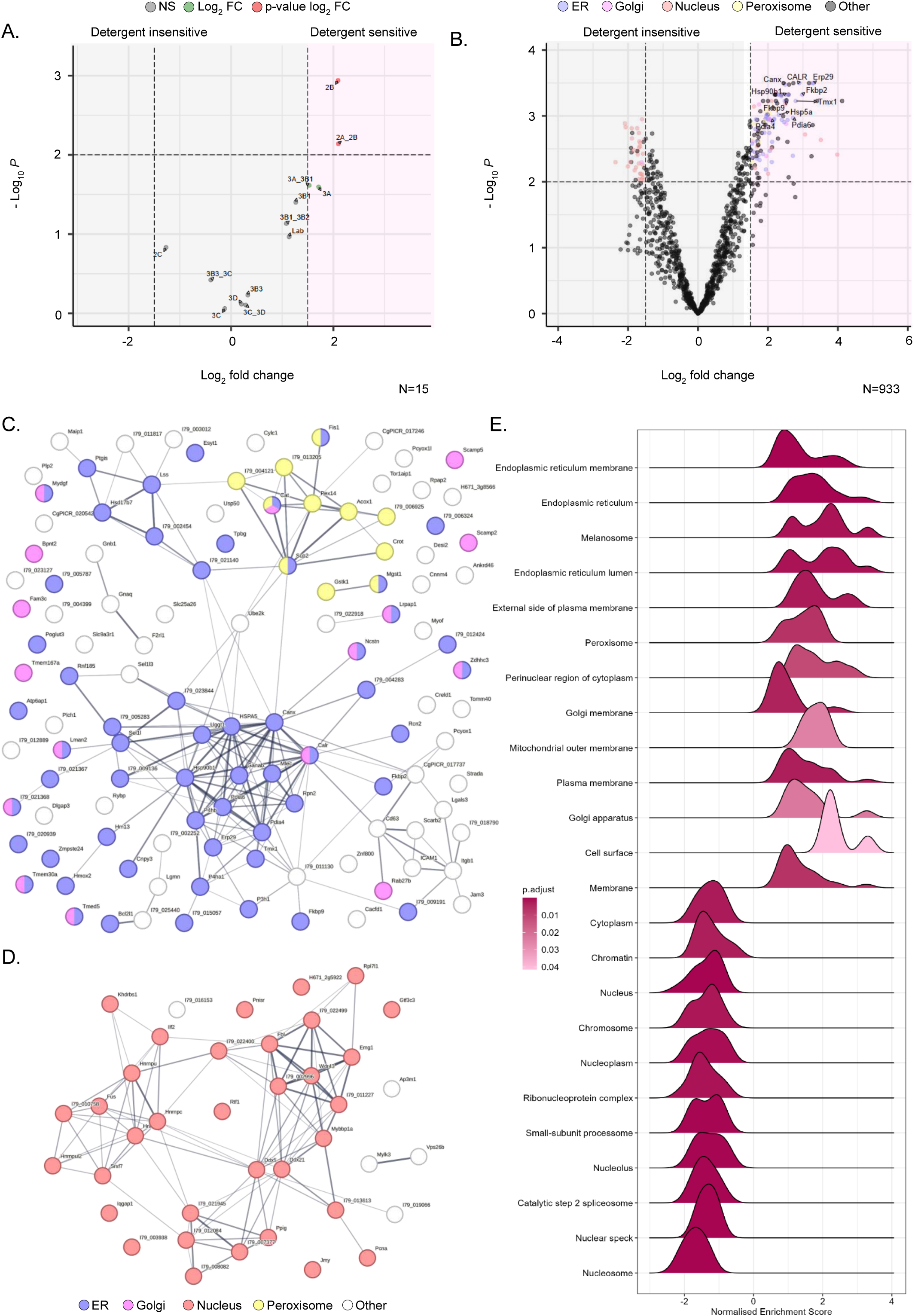
TMT mass spectrometry analysis of 2C co-immunoprecipitated proteins in the absence and presence of detergent. (A) BHK-21 cells transfected with wild-type FMDV replicon were homogenised 4-hpt and either detergent-treated or non-treated before anti-2C immunoprecipitation. Co-immunoprecipitated proteins were identified by TMT mass spectrometry and peptides mapped against the FMDV replicon polyprotein. Relative log_2_ fold change of each protein abundance was plotted comparing non-treated:detergent-treated. (B) Peptides mapped to host proteins were analysed and significant hits were colour coded by cellular compartment. STRING analysis was performed on host protein hits >1.5 log_2_ FC & <0.01 adj. p-value (C) or <1.5 log_2_ FC & 0.01 adj. p-value (D). Protein nodes are colour coded based on cellular compartment (E) Gene enrichment analysis was performed comparing non-treated and detergent-treated conditions. Relative enrichment of Gene ontology terms is displayed in a ridgeline plot, +ve values represent enrichment. Data were collected from three biological repeats.

Having established that the differential enrichment analysis of viral proteins followed the expected trend, we investigated host protein changes following detergent treatment. Plotting these changes and colour coding significantly enriched proteins (≥ 1.5 or ≤ -1.5 log_2_ FC & ≤ 0.01 adjusted p-value) for their respective cellular component Gene Ontology (GO) term, a clear enrichment of proteins associated with the endoplasmic reticulum (ER) (GO:0005783) was evident (Figure 5.B). These accounted for 57 out of a total of 118 proteins significantly enriched and detergent sensitive. Other membranous compartments were represented in this group, including 16 Golgi-associated (GO:0005794) and 11 peroxisome-associated (GO:0005777) proteins. To visualise the physical and function relationships between these 118 significantly enriched detergent sensitive proteins, STRING analysis was performed (Figure 5.C). Clustering of ER- and peroxisome-associated proteins was evident, including the ER proteins calnexin, calreticulin, Pdia4 & 6, HSP5 & 90, Erp29 and Tmx1. Conversely, there appeared to be no clear clustering evident for Golgi-associated proteins in the STRING analysis.

Similar analysis was conducted to the subset of proteins which displayed a significant negative enrichment (≤-1.5 log_2_ FC & 0.01 adjusted p-value), likely indicative of their detergent-insensitivity (Figure 5.B). These proteins numbered 39 with 34 associated with the nucleus (GO:0005634) and roughly clustering by STRING analysis into two interlinked groups, one containing a number of RNA splicing related proteins such as HNRNPs and a second containing ribosome biogenesis associated proteins such as Emg1 (Figure 5.D).

To further investigate the cellular component composition of the immunoprecipitated proteins, we performed gene set enrichment analysis (GSEA) plotting cellular component for non-treated compared to detergent-treated (Figure 5.E). Visualisation of the data as a ridgeline plot highlighted that some terms within the dataset contained subpopulations of greater enrichment (two or more peaks) compared to those with a global enrichment (single peak). As expected, the normalised enrichment scores (NES), show an overrepresentation of membrane-associated terms and an underrepresentation of nuclear terms in agreement with the previous differential enrichment analysis. Further to this, terms related to the ER predominate in the GSEA with other compartments previously mentioned, Golgi and peroxisome are also represented. Additionally, several other terms such as the melanosome and the cell membrane were also highlighted to be enriched, which was not predicted or easily discernible from the DEA volcano plot and STRING analysis. Perhaps less expected are the bimodal distributions observed for some terms, for example ER membrane and cell surface, while most nuclear-related terms display a single peak suggesting a more homogenous enrichment. Taken as a whole, the analysis of the enriched proteins following anti-2C immunoprecipitation in the presence or absence of detergent highlighted a predominance of proteins related or associated with the ER as well as the detergent insensitivity of a number of nuclear proteins involved in RNA biology.

## Discussion

The data presented here demonstrate interactions between viral proteins 2C, 3A (and precursors), 3CD and a number of host proteins. Evidence for the interaction between 2C and 3A being membrane-mediated was provided through sensitivity to detergent. The membrane associated nature of 2C was further highlighted with the identification of 57 significantly enriched ER related proteins. These results support a proposed model of predominantly ER-derived membranous FMDV replication complexes containing 2B, 2C, 3A along with more transient interactions with 3CD and 3D.

Several studies have taken a reductionist approach to investigating the role of 2C in FMDV replication (Moffat *et al*., 2005; Knox *et al*., 2005; Moffat *et al*., 2007; Sweeney *et al*., 2010; Mahajan *et al*., 2021; Zhang *et al*., 2022). Transfection of 2BC and 2B/2C alone into cells has demonstrated the membrane re-modelling ability of both proteins, mimicking the replication complexes formed during infection (Moffat *et al*., 2005; Moffat *et al*., 2007). Interestingly, this approach also found that 2C alone (or precursor 2BC) was unable to halt ER-Golgi transport (Moffat *et al*., 2007). The data presented here, while also adopting a reductionist approach, sought to identify the interactions by using an FMDV replicon model, focusing on 2C as a proxy for the replication complex.

The relationship between 2C and 3A precursors was investigated here by different treatments of transfected lysates. Through this approach, it was found that the interaction between these proteins was likely membrane-mediated and could be disrupted by >0.1% Triton X-100, yet was tolerant to high salt concentration, suggesting that this interaction was not electrostatic (Dumetz *et al*., 2007). While a robust interaction between 2C and 3A precursors was observed, that between 2C and 3CD and 3D was less strong. This suggests that the replication complex of FMDV has some similarity to that of PV, with 2C and 3A directly associated with a membrane alongside interactions with other proteins such as 3CD and 3D (Barton, Black and Flanegan, 1995).

After demonstrating co-immunoprecipitation of viral proteins through western blotting, we interrogated the detergent-sensitive interacting partners of 2C using mass spectrometry. By mapping viral peptides to their respective proteins, we reconfirmed our previous observations that the co-immunoprecipitation of 2C and 3A/3A precursors was disrupted through detergent treatment, providing a validation for our subsequent analysis. Peptides mapping to 2B were shown to be co-immunoprecipitated with 2C in a detergent-sensitive manner, similar to that of 3A. We had not seen this interaction previously due to the lack of antibody reagents to 2B. Due to the change in 2B peptide detection between the two treatment conditions we can propose that the association of 2B and 2C occurred predominantly as individual proteins rather than 2BC. The ability of 2BC (and a synergistic ability of co-expressed 2B and 2C) to block ER-Golgi trafficking has been reported previously (Moffat *et al*. 2005; Moffat *et al*. 2006). While some precursors may have been present across both sample sets, the large amount mature 2B associated with these putative replication complexes is perhaps further evidence of the co-operative nature of 2B and 2C proteins when associated with membranes during FMDV replication. While 3D was detected in both conditions, the relative ratio of 3D was not significantly affected by detergent treatment, suggesting that its interactions with 2C, and possibly by extension with the replication complex, is more dependent on protein-protein or protein-RNA interactions. This would be in-line with data reported on the multimerisation of 3D, its ability to function *in vitro* in the absence of membranes, and its association with 3A precursors (Bentham *et al*. 2012; Loundras *et al*. 2022; Ferrer-Orta, Ferrero and Verdaguer, 2023).

Having demonstrated that viral proteins interacting with 2C were enriched or depleted as expected following detergent treatment, we extended the analyses to include host proteins. The composition of FMDV replication complexes has remained elusive with contention over the origins of the membranes involved as well as the content of these structures. Through our analysis of peptides detected following 2C immunoprecipitation in the presence or absence of detergent treatment, we have identified an enrichment of ER-associated proteins, suggesting this could be the predominant source of replication complex membranes. The cellular component GO term for the ER analysis identified 57 member proteins >1.5 log_2_ fold. These enriched proteins were split between the daughter terms for both the organelle membrane and lumen, supporting the membrane-associated nature of these interaction with 2C and the replication complex. This is in contrast to the lack of Golgi-associated terms, which were far less enriched.

Previous studies using ER and Golgi marker proteins have not identified a robust association with either 2C or 3A of FMDV (Knox *et al*. 2005; Garcia-Briones *et* al. 2006, Berryman *et al*. 2012; Midgley *et al*. 2013). A previous attempt to identify membrane components co-immunoprecipitating with 2C was unable to positively blot for ER or Golgi markers (Knox et al. 2005). Here, we present supportive data i.e. the absence of GM-130 (Golgi marker) TMT, which is traditionally the protein of choice for these compartments. Instead, our data suggest that 2C associates with an array of other ER proteins, including calnexin, and a relative lack of Golgi proteins. Additionally, a small number of peroxisome-associated proteins were also identified as detergent-sensitive. The functions of these remains to be investigated in the context of FMDV replication, however, there is growing evidence of the involvement of peroxisomes in the replication of other RNA viruses (Ferreira *et al*., 2022). The wider list of detergent-sensitive 2C interactors, beyond calnexin, provides a number of future targets for visualisation, characterisation and potential disruption of the establishment or function of the FMDV replication complex.

Our mass spectrometry analysis also suggested a number of nuclear proteins immunoprecipitated with 2C in a detergent-insensitive manner. A large proportion of these proteins are canonically involved in RNA processing activities with many being splicing factors such as serine/arginine splicing factor 7 (SRSF7). There is emerging evidence across both DNA and RNA viruses for the ability to modify nuclear splicing machinery, suggesting a potentially important requirement for these functions in viral replication (Harper *et al*., 2024; Castello and Kamel, 2025). Experimental data using human rhinovirus identified a number of nuclear splicing factors which migrated from their canonical localisation into the cytoplasm during infection (Flather *et al*., 2018). Additionally, degradation of the nuclear lamina and disruption of nuclear pores have also been previously identified to occur during FMDV replication. Our data support these observations and suggest that nuclear proteins may interact with 2C in a protein-protein or protein-RNA-dependent manner in the replication complex. The function and redundancy of these nuclear proteins during FMDV replication is an area requiring further investigation yet may yield potential targets for the disruption of viral replication machinery.

Probing with anti-2C, anti-3A and anti-3D antibodies in western blots produced from primary and secondary gradients displayed an enrichment in ∼20% iodixanol. This enrichment was maintained following secondary gradient purification and is similar to results described by Pietilä *et al*. when purifying replication complexes from SFV infected cell lysates (Pietilä, van Hemert and Ahola, 2018). The co-sedimentation of multiple FMDV proteins seen here, with different molecular weights ranging from ∼20 to ∼75 kDa suggests their presence as part of a complex. Blocking host-cell transcription through treatment with actinomycin D followed by supplying tritiated uridine to cells transfected with either wild-type or GNN replication-deficient replicon enabled detection of vRNA. Detection of [^3^H] was at maximum in the same fractions enriched for viral proteins. In this experiment total vRNA is likely to have been labelled, i.e. both +ve sense and -ve sense species. Future work to further characterise which types of viral RNA were co-sedimenting could employ a strand-specific RT-qPCR approach which can differentiate +ve and -ve sense viral RNA extracted from FMDV-transfected cells (Dobson *et al*., 2023). Taken together, co-sedimentation of protein and vRNA suggests that these are inter-related, lending support to an enrichment of putative replication complexes.

Overall, the data presented here supports a model of FMDV replication complexes derived predominantly from membranes of ER origin, containing at least 2B, 2C, various precursor forms of 3A, 3CD and 3D, as well as a range of host proteins.

## Acknowledgments

N.J.S, M.R.H, D.J.R and S.J.D were funded by Biological Sciences Research Council (BBSRC) of the United Kingdom (research grant BB/K003801/1); E.V.H, N.J.S and D.J.R received funding from the US National Institutes of Health (NIH Grant R01 AI 16945). M.R.H. received funding from the Medical Research Council of the United Kingdom (research grant MRC (MR/S007229/1); J.F. received funding from PID2023-149259NB-I00, MICIU/AEI/ 10.13039/501100011033 and “ERDF A way of making Europe”; CH was funded by a BBSRC studentship (BB/M011151/1). We thank Francisco Sobrino (Centro De Biologia Molecular Severo Ochoa, Madrid) for the gift of FMDV primary antibodies. M.R.H., N.J.S., J.F. and D.J.R. designed the study; C.H, MRH and S.J.D. conducted experimental work; C.H., and E.V.H. analyzed data; M.R.H., N.J.S., J.F. and D.J.R. provided supervision. C.H. prepared the draft manuscript and all authors contributed to the final version. The funders had no role in the study design, data collection and analysis, decision to publish or preparation of the manuscript.

