## Supplementary data for "Interactions between the 2C protein of FMDV and components of the viral replication machinery are mediated by ER-derived membranes"

### Supplementary information

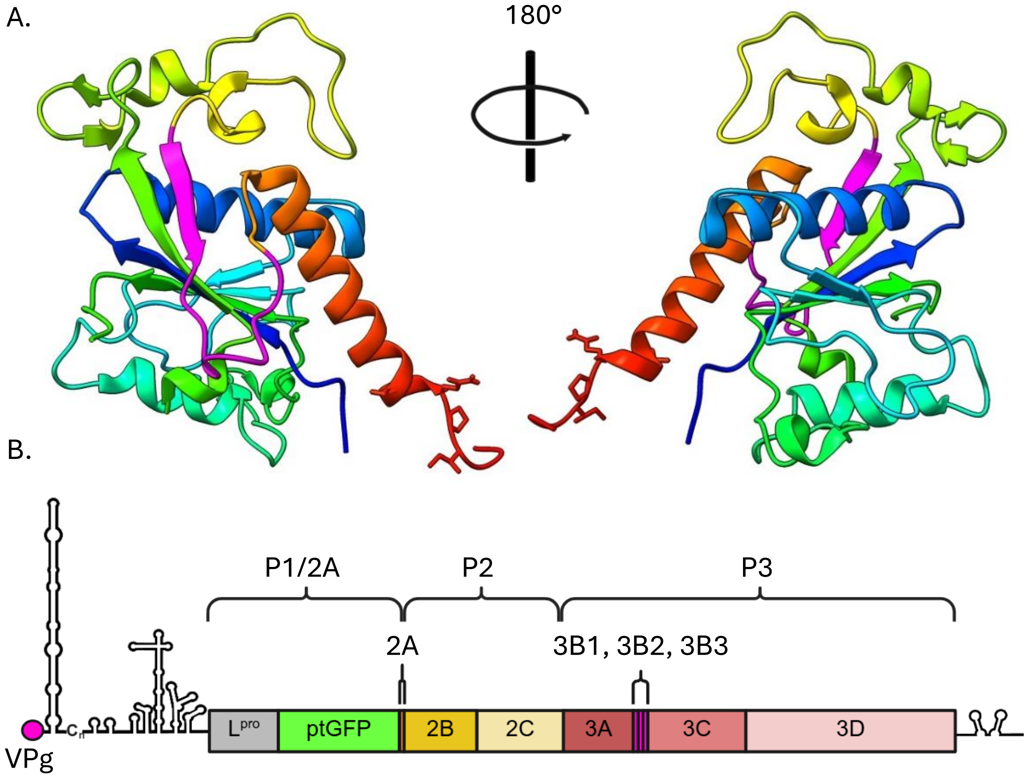


Figure S1. (A) Peptide residues (pink) used to generate novel anti-2C antibody mapped onto the crystal structure of FMDV protein residues 97-318 aa (PDB 7E6V), coloured N- to C- terminal rainbow gradient (Zhang *et al.*, 2022).

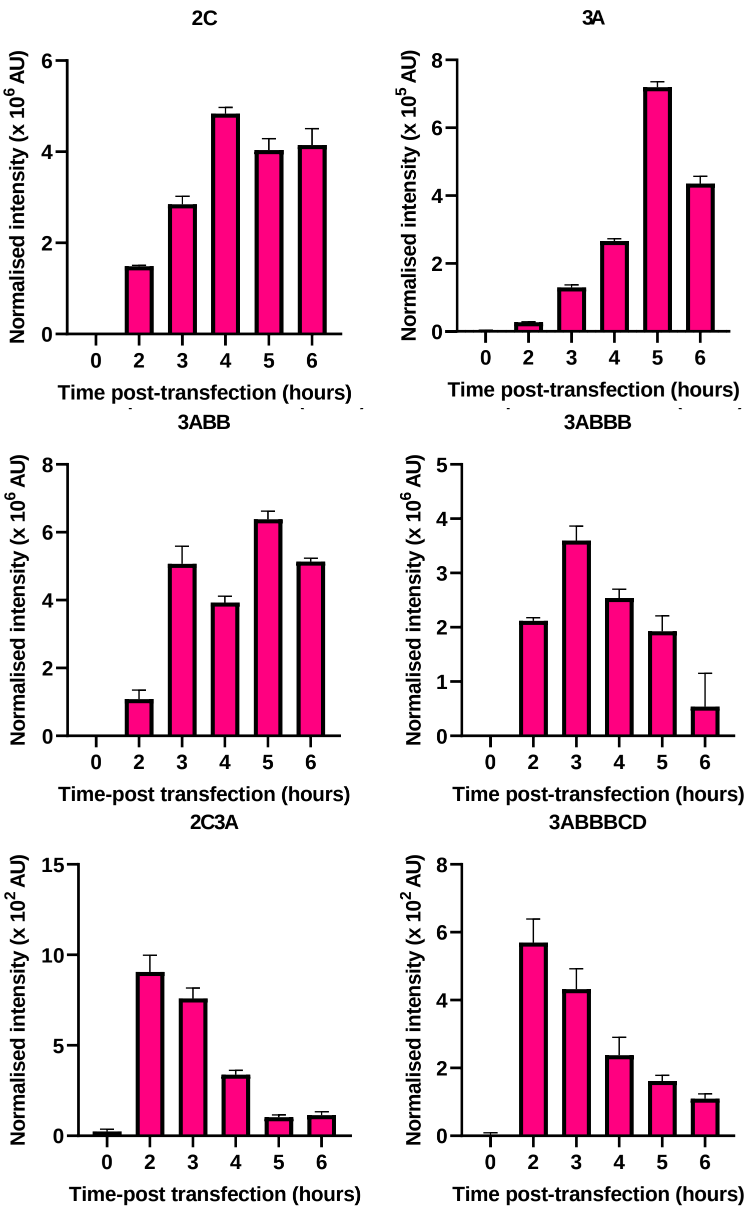


Figure S2. Densitometry of FMDV protein expression over time.

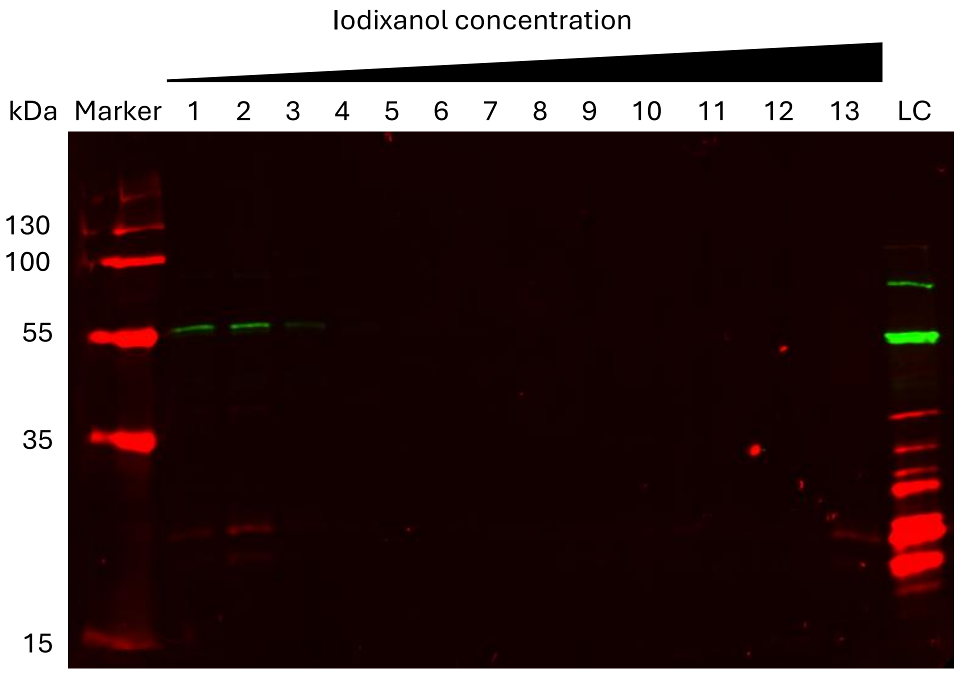


Figure S3. Primary gradient purification of GNN transfected lysate. Probed with anti-3D and anti-3A. LC = Loading control of wild-type transfected lysate.

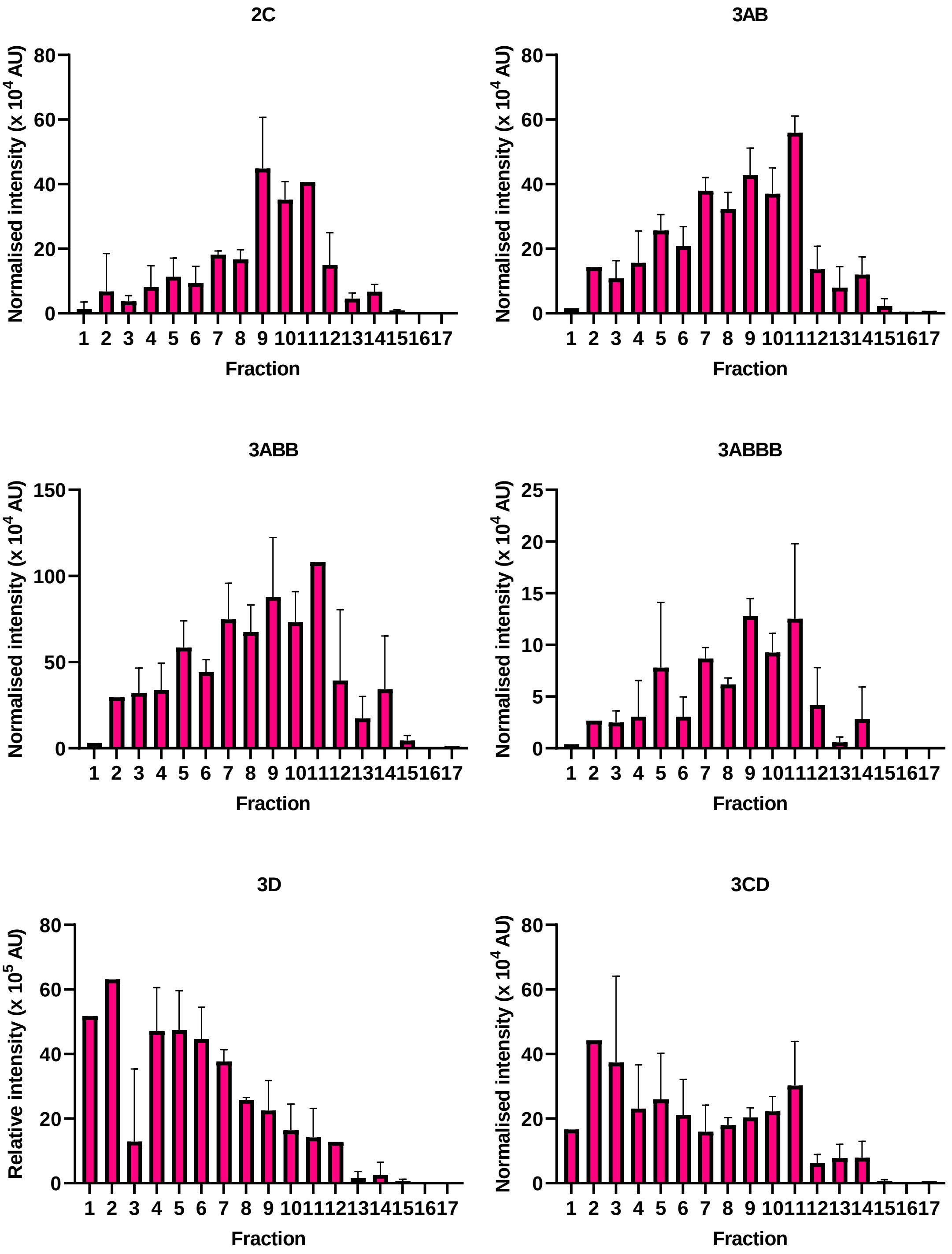


Figure S4. Densitometry of FMDV protein signal across a primary gradient purification. Normalised to loading control of wild-type transfected lysate.

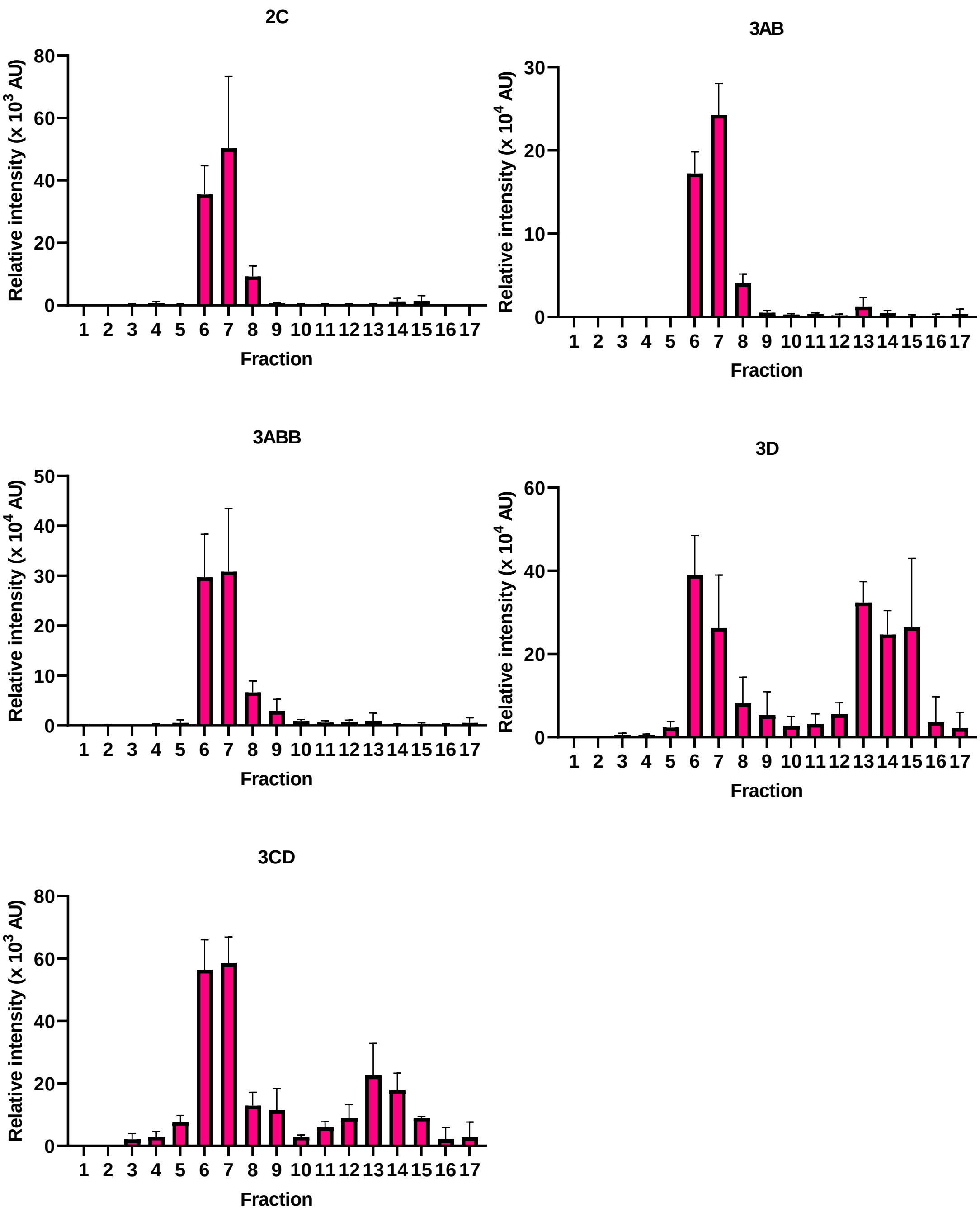


Figure S5. Densitometry of FMDV protein signal across secondary gradient purification. Normalised to loading control of wild-type transfected lysate.
